# Conditional distribution modeling as an alternative method for covariates simulation: comparison with joint multivariate normal and bootstrap techniques

**DOI:** 10.1101/2020.11.01.363325

**Authors:** Giovanni Smania, E. Niclas Jonsson

## Abstract

Clinical trial simulation (CTS) is a valuable tool in drug development. To obtain realistic scenarios, the subjects included in the CTS must be representative of the target population. Common ways of generating virtual subjects are based upon bootstrap (BS) procedures or multivariate normal distributions (MVND). Here, we investigated the performance of an alternative method based on conditional distributions (CD). Covariates data from a hypertension drug development program were used. The methods were evaluated based on the original dataset (internal evaluation) and on their ability to reproduce an older, unobserved population (extrapolation). Similar results were obtained in the internal evaluation for summary statistics, yet BS was able to preserve the correlation structure of the empirical distribution, which was not adequately reproduced by MVND; CD was in between BS and MVND. BS does not allow to extrapolate to an unobserved population. When the dataset used to inform the extrapolation was well approximated by a MVND, the results from CD and MVND were comparable. However, improved extrapolation performance was observed for CD when deviations from normality assumptions occurred. If CTS is used to simulate within the observed distribution, BS is the preferred method. When extrapolating to new populations, a parametric method like CD/MVND is needed. In case the empirical multivariate distribution is characterized by linearly related covariates and unimodal marginal distributions, MVND can be used because of the simpler statistical framework and well-established use; however, if uncertainty about the MVND assumptions exists, CD will increase the confidence in the simulations compared to MVND.

## Introduction

In several research fields that aim to study complex systems, mathematical modeling is routinely used to quantitatively describe the system, generate or test hypotheses and explore system behavior by means of model simulations. Drug discovery and development is one such field, that has seen the application of modeling and simulation - or pharmacometrics - growing at a fast pace in the last decades [1]. Arguably, the success of pharmacometrics lies in its ability to integrate and synthesize evidence collected from heterogenous sources throughout the drug development spectrum, ultimately providing a sound quantitative input to the decision-making process in pharmaceutical R&D [2-4].

In the pharmacometrician’s toolkit, clinical trial simulation (CTS) offers the possibility to mitigate the risk of study failure by prospectively exploring the performance of a given study design and/or of the decision-making criteria [5]. Among others, applications of CTS include proof-of-concept study design [6], outcome prediction in phase 3 [7], pediatric trial optimization [8,9], evaluation of non-adherence [10,11] and guidance in dose titration decisions [12]. CTS can be broken down into three building blocks: a disease model, a drug model and a trial model [13]. The trial model requires the definition of a patient population through a set of characteristics, often termed covariates; in order to obtain realistic simulation scenarios, the set of covariates included in the CTS must be representative of the target population.

Common ways of generating covariate distributions for CTS are based upon non-parametric bootstrap (BS) procedures or multivariate normal distributions (MVND) [14,15]. While BS is a non-parametric method, MVND requires certain parametric assumptions to be met by the original data set at hand. The different nature of these two methods brings respective pros and cons: with BS it is not possible to extrapolate outside of the observed distribution, but it preserves the relationships between covariates, thereby returning simulated subjects that are physiologically plausible. Conversely, the parametric distribution underlying the MVND -under certain assumptions about the means and correlations for the simulated population - can be used to generate completely new individuals. However, even with appropriate assumptions, the unrestricted nature of the underlying multivariate distribution may still lead to unrealistic simulated subjects.

Multiple imputation [16,17] is a technique used to handle complex missing data problems in statistical analysis, where multiple imputed data sets are analyzed separately, and inference is made on the pooled results. Missing data are imputed by sampling from conditional distributions using a pre-specified imputation model [18]. A commonly used imputation model is predictive mean matching (PMM) [19]. For each missing entry, PMM forms a small set of candidate donors from all complete cases that have predicted values closest to the predicted value for the missing entry. One donor is randomly drawn from the candidates, and the observed value of the donor is taken to replace the missing value. It follows that missing data are imputed with real values observed elsewhere in the data set, so imputations outside the observed data range will not occur. This feature makes sampling from conditional distributions using PMM (CD) an appealing method for covariate simulations, since in principle it embraces the advantages of MVND and BS, i.e., it can retain physiological plausibility while allowing to extrapolate outside of the observed multivariate distribution.

The objectives of this analysis were to investigate the operating characteristics of CD when used to simulate covariates distributions, and to compare them with those of BS and MVND.

## Methods

### Data

The data set used to build the simulations, hereafter referred to as the original data set, was obtained from a hypertension drug development program with data in 233 healthy subjects (HS) and 706 patients. The original data set contained the baseline values of the following covariates: age, weight (WT), serum creatinine (SCR), creatinine clearance (CRCL), sex and race. The summary statistics of the covariates in the original data set are reported in Table 1. The histograms of the continuous covariates stratified by HS vs. patients and the scatterplot matrix of the log-transformed continuous covariates are shown in Figure 1A and 1B, respectively.

**Table 1.** Summary statistics of the covariates in the original data set.

|  |  |
| --- | --- |
| Total number of subjects | 939 |
| **Continuous covariates** |  |
| Age (years) |  |
| Mean | 46.4 |
| SD | 12.4 |
| Median | 47.0 |
| Range | 18.0-77.0 |
| Weight (kg) |  |
| Mean | 88.5 |
| SD | 20.1 |
| Median | 86.6 |
| Range | 46.3-172 |
| Serum creatinine (μmol/L) |  |
| Mean | 78.5 |
| SD | 16.2 |
| Median | 78.7 |
| Range | 41.5-133 |
| Creatinine clearance (mL/min) |  |
| Mean | 124 |
| SD | 34.3 |
| Median | 120 |
| Range | 47.0-282 |
| **Categorical covariates** |  |
| Population, n (%) |  |
| Patients | 706 (75.2) |
| Healthy subjects | 233 (24.8) |
| Sex, n (%) |  |
| Males | 534 (56.9) |
| Females | 405 (43.1) |
| Race, n (%) |  |
| White | 737 (78.5) |
| Black | 179 (19.1) |
| Asian | 19 (2.0) |
| American Indian | 2 (0.2) |
| Other | 2 (0.2) |

**Figure 1.**
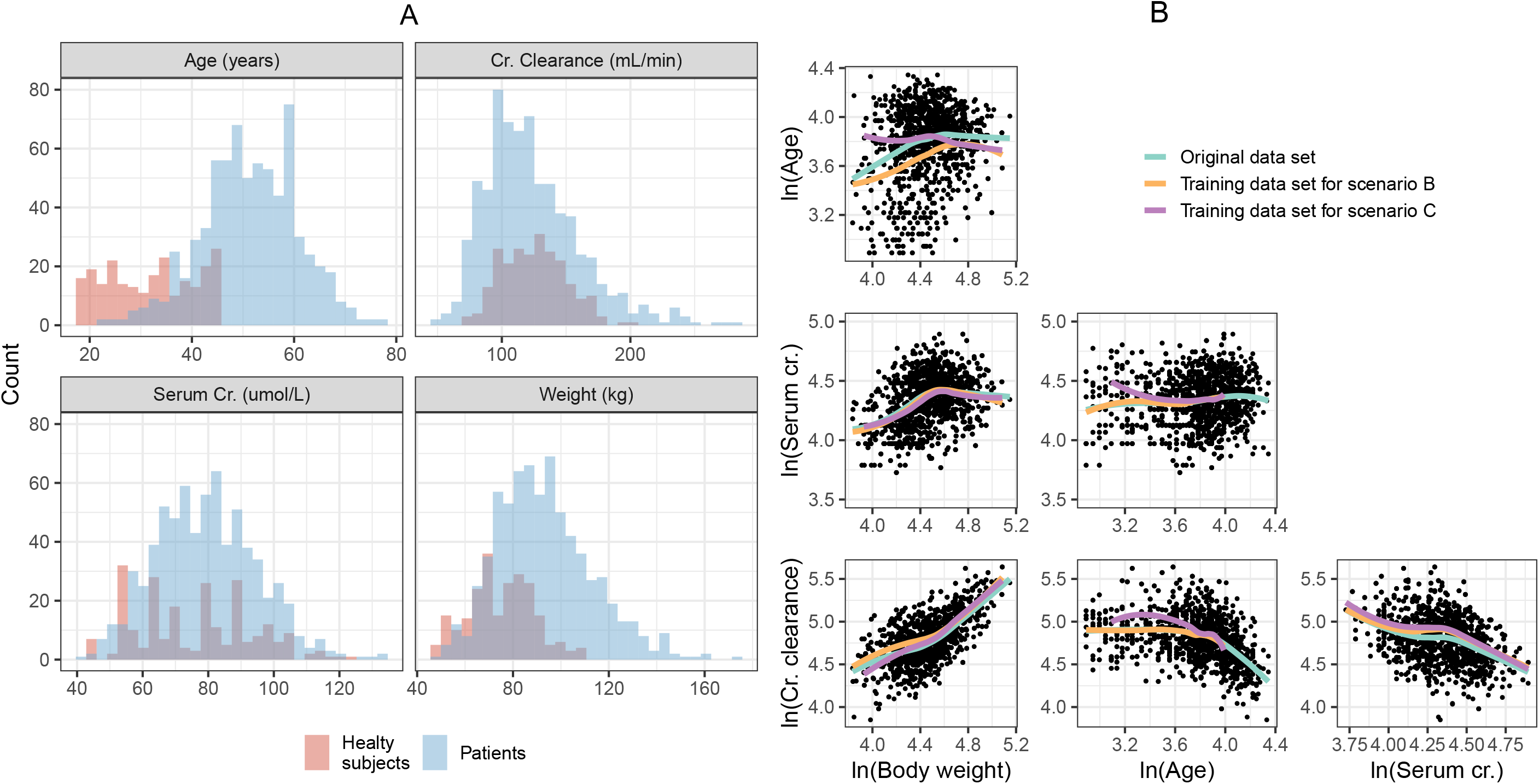
Histograms of continuous covariates colored by patients vs. healthy subjects (A) and correlation between pairs of continuous covariates (B). The solid line is a loess regressor.

### Covariate simulation methods

#### BS

The BS method consisted of sampling, with replacement, covariate vectors from the original data set.

#### MVND

The MVND was implemented based on the framework presented in Tannenbaum et al. [15], that is briefly described. Before estimating the parameters of the MVND from the original data set, continuous covariates are log-transformed to constrain them to be positive; the covariate vectors simulated with the MVND are then exponentiated back to obtain the actual covariate values. As to categorical covariates, no pre-transformation is used, and they are treated as if they were continuous variables. This implies that the MVND can generate non-discrete values for categorical covariates, which then must be mapped back to their respective category based on a continuous critical value (see [15] for more details). Two important assumptions of the MVND are that the covariates in the original data set follow the same known distribution, and that they are linearly related to each other. A third assumption is that all the marginal distributions should not display clear multimodal trends. No truncation was applied to the covariates simulated with the MVND.

#### CD

Simulations with CD were performed using the fully conditional specification (FCS) algorithm (also known as multivariate imputation by chained equations) as implemented in the R package *mice* version 3.6.0 [19]. FCS is an iterative method in which missing data are imputed on a variable-by-variable basis. Let the covariate data set be represented by the matrix Y, FCS specifies a group of conditional densities P(Y_j_^mis^ | Y_j_^obs^, Y_-j_,>,□), where Y_j_^mis^ and Y_j_ ^obs^ are the set of missing and observed values for the j-th covariate, respectively, Y_-j_ are the values of all covariates in Y that are not the j-th one, and □ is the imputation model. Starting from simple random draws from the marginal distribution, imputation under FCS is done by iterating over the conditionally specified imputation models (an illustrative example using standard linear regression as imputation model is shown in Figure S1). The complete FCS algorithm as implemented in the *mice* package is reported in Supplementary Information 1, further details can be found in [16,18].

The *mice* package offers the opportunity to choose from a selection of linear imputation models; here, the default methods were used, namely PMM [19] and multinomial logistic regression for continuous and categorical covariates, respectively. PMM first calculates the predicted value of the target covariate according to the specified linear regression model, secondly forms a set of donors/neighbors from the original data set constituted by the K closest values to the predicted one (K=5 in the present analysis), and finally makes the actual prediction by randomly drawing one value form the set of donors. Thus, the value of the simulated covariate is ultimately drawn from the original data set.

The CD simulations were performed as imputation of data sets where all covariates were assumed to be missing, leveraging the original data to train each univariate imputation model (Figure S1). By doing so, the probability of a certain covariate being missing does not depend on the unobserved values of that (or other) covariate(s), that is, the missing covariates are said to be missing at random (MAR); this in turn implies that the missing data mechanism is ignorable, a condition that can be handled by the FCS algorithm [16,18]. An ad-hoc R function was developed and used to automate covariates simulation with CD (Supplementary Information 2).

### Simulations set up and methods comparison

N=30 replicates of the same size as the original data set were simulated using BS, MVND and CD. CRCL was not directly simulated but derived from the simulated values of WT, SCR and sex using the Cockcroft-Gault [20] formula, similarly to what was done in the original data set.

In a first step, mean, standard deviation (SD), median, range and variance-covariance matrix for continuous covariates and proportions for categorical covariates were calculated for each of the 30 simulated replicates. Secondly, for each replicate *r*, the relative prediction error between the statistic of the simulated data set (P_sim,r_) and the original statistic (P_org_) was computed as rPE_r_ (%) = (P_sim,r_ -P_org_)/P_org_*100. Finally, relative bias (rBias) and root mean squared error (rRMSE) were derived as 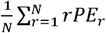 and 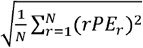, respectively.

The comparisons described above were carried out under three different scenarios. Scenario A used the entire original data set to inform the BS, MVND and CD simulations (internal evaluation). In Scenario B, with the aim of investigating the extrapolation performance of the methods, the original data set was divided into a training and a test data set, where only the training data set was used to inform the simulations, while the test data set was used to assess the predictive performance of the methods (external evaluation, P_org_ in this case is the statistics computed in the test data set). The training data set was defined as all subjects in the original data set younger than 55 years of age, mimicking a CTS exercise executed to investigate the features of a clinical trial performed in an older, unobserved population. Scenario C was the same as Scenario B but excluding HS from the training data set (the histograms of the continuous covariates stratified by sex in the training and test data sets for Scenarios B and C are displayed in Figure S2). By definition, the BS does not allow to extrapolate outside of the empirical distribution, hence Scenarios B and C were tested for CD and MVND only. In Scenario B and C, the MVND simulations for age were obtained by truncating the distribution in the range 55-77 years. The age values simulated by the MVND were then used to seed the CD simulations, that is, the age values were not imputed but assumed to be known (see example C in Supplementary Information 2). Accordingly, only the covariance terms for age were included in the evaluation, whereas the other age statistics were not. It should be noted that in the CD simulations under Scenarios B and C the MAR assumption still holds true, as the probability of the missing data depends on the observed data, i.e., the probability of a covariate being missing is higher in older subjects compared to younger ones.

## Results

### Scenario A

Accuracy (rBias) and precision (rRMSE) of the continuous statistics are shown in Figure 2A-B. The mean and median of the original data set were maintained in the simulated population for all the methods. Likewise, SD and range were well estimated by BS and CD, whereas MVND resulted in larger rBias and rRMSE for the range of all covariates and for the SD of age. Figure 2C shows that the overall operating characteristics for the simulation of categorical covariates were similar across the three methods. BS was able to preserve the correlation structure of the empirical distribution, which was instead not adequately reproduced by MVND; CD was in between BS and MVND (Figure 3A). This in turn resulted in the lowest (BS) and highest (MVND) rBias and rRMSE in the elements of the variance-covariance matrix (Figure 3B-C).

**Figure 2.**
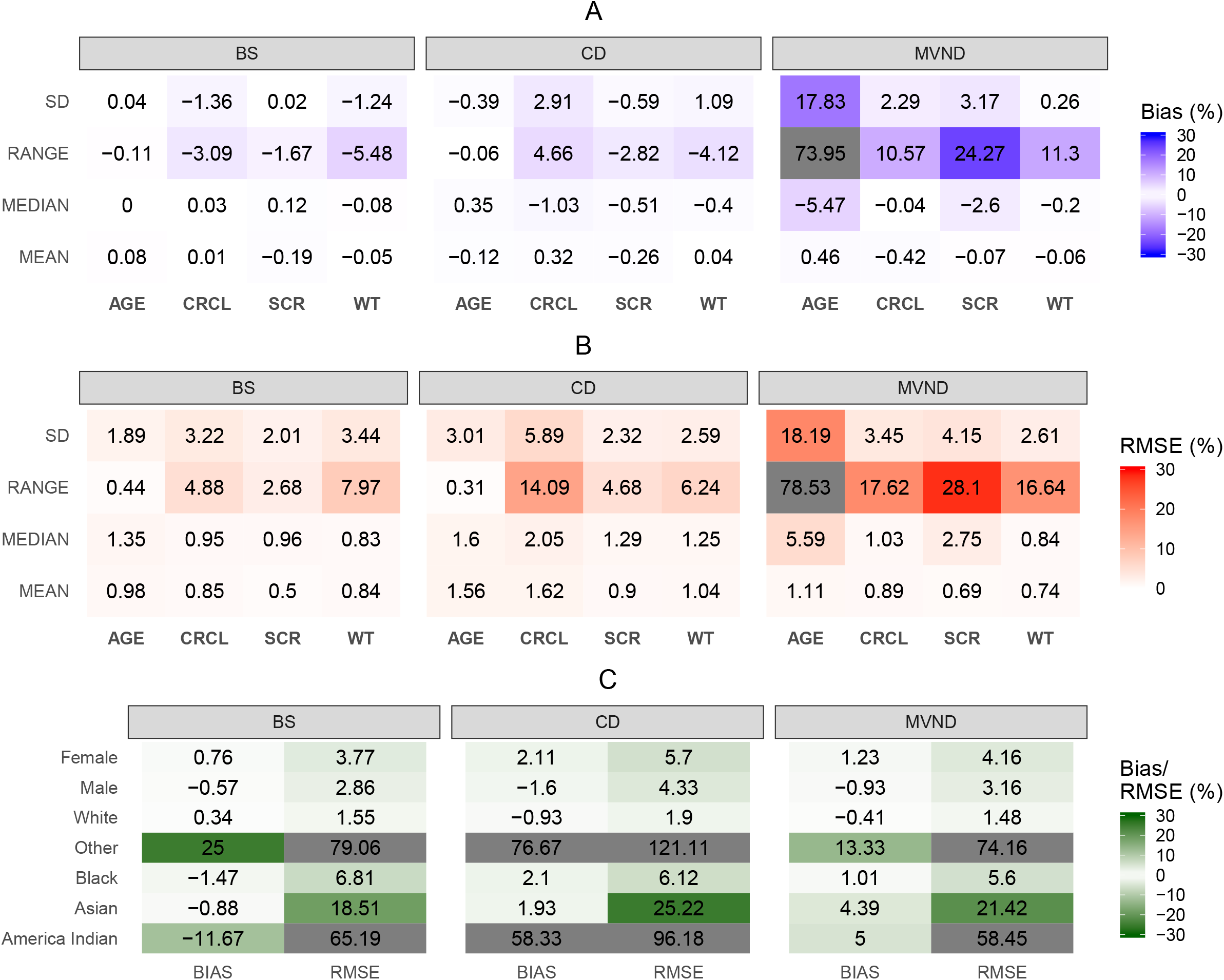
Relative Bias and RMSE for summary statistics of continuous (A and B) and categorical (C) covariates in Scenario A (cells where the absolute value of relative Bias/RMSE is greater than 30% are grayed out).

**Figure 3.**
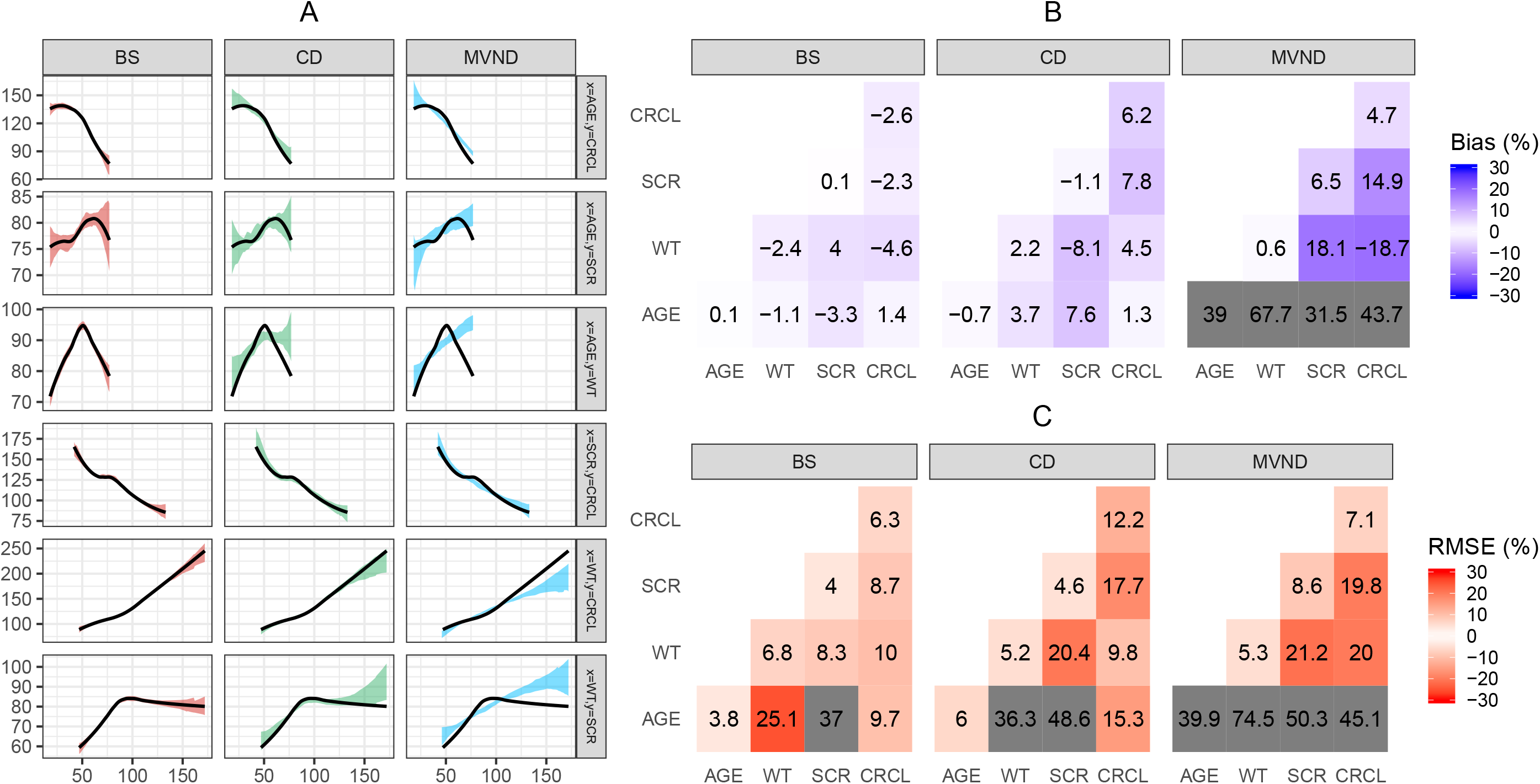
Visual predictive check of the relationship between pairs of continuous covariates (A): the black line represents a loess regressor through the original data set, while the shaded area depicts the 80% confidence interval of the loess regressor fitted to each of the 30 replicates. Relative Bias (B) and RMSE (C) for the variance-covariance matrix of the continuous covariates (cells where the absolute value of relative Bias/RMSE is greater than 30% are grayed out). Scenario A.

### Scenario B

Using CD for the simulation of an older patient population led to lower rBias and rRMSE in the summary statistics of the continuous covariates, compared to MVND (Figure 4A-B). Figure 4C shows that the prediction of the proportion of males and females in the extrapolated data set was more accurate and precise for MVND vs. CD, while similar performance was obtained for race.

**Figure 4.**
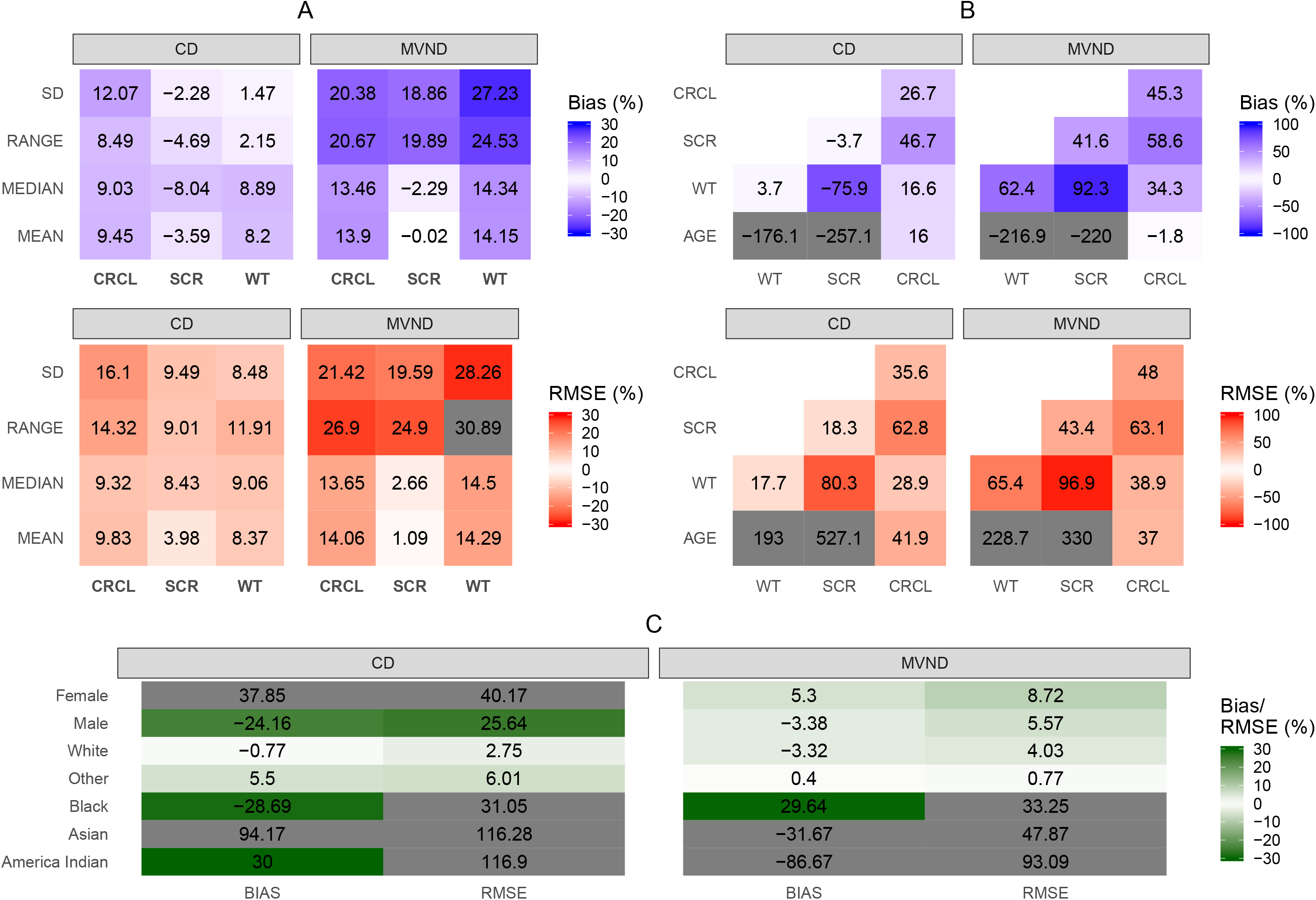
Relative Bias and RMSE for summary statistics of continuous (A) and categorical (C) covariates in Scenario B (cells where the absolute value of relative Bias/RMSE is greater than 30% are grayed out). Relative Bias and RMSE for the variance-covariance matrix of the continuous covariates (B) in Scenario B (cells where the absolute value of relative Bias/RMSE is greater than 100% are grayed out). There were no subjects with “race = other” in the training data set, therefore in subfigure C rBias and rRMSE were computed using the absolute difference instead of the relative one for this category.

### Scenario C

When the HS data were removed from the training data set, no clear differences were observed between CD and MVND: some continuous covariates were better predicted by MVND (e.g. serum creatinine) and some others by CD (e.g. WT) (Figure 5A-B). As displayed in Figure 5C, sex was still more accurate and precise for MVND vs. CD.

**Figure 5.**
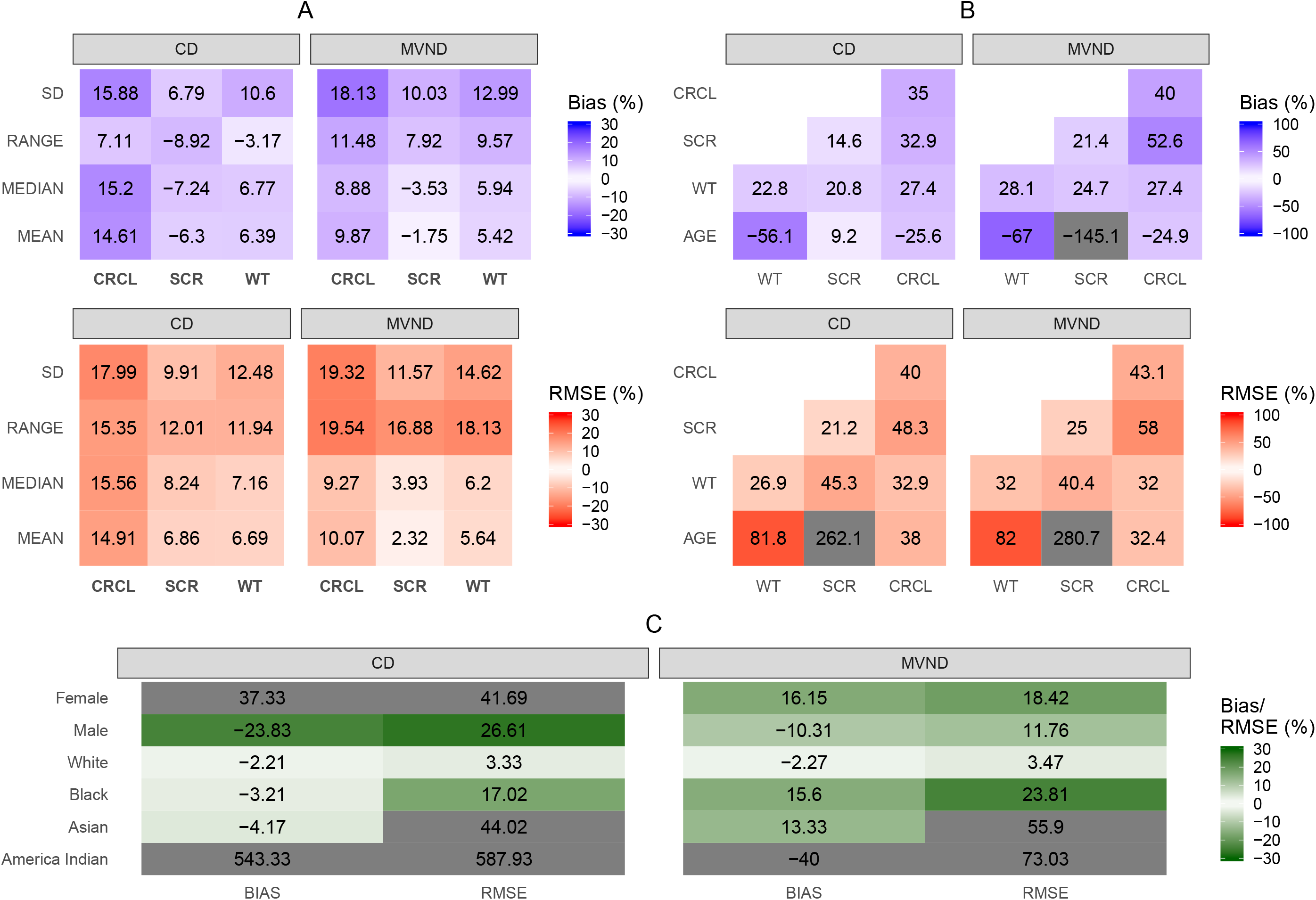
Relative Bias and RMSE for summary statistics of continuous (A) and categorical (C) covariates in Scenario C (cells where the absolute value of relative Bias/RMSE is greater than 30% are grayed out). Relative Bias and RMSE for the variance-covariance matrix of the continuous covariates (B) in Scenario C (cells where the absolute value of relative Bias/RMSE is greater than 100% are grayed out).

## Discussion

To maximize the predictability of a CTS exercise, plausible covariate values -representative of the target patient population - must be included in the trial model. Previous studies have investigated the operating characteristics of MVND as a tool to generate virtual populations [15], and compared them with BS approaches [14]. In the present work, we introduced an alternative approach to covariate simulations that borrowed the methodology used for multiple imputation of incomplete data sets in statistical analysis [16,17]. In addition, we have also explored the extrapolation performance of the methods in predicting the summary statistics of an older, unobserved patient population.

In the internal evaluation (Scenario A) the three methods adequately reproduced the summary statistics of the continuous covariates present in the original data set (Figure 2A-B). The results indicated that, for some covariates like age and SCR, dispersion parameters were not well predicted by MVND, which in turn led to the generation of implausible virtual subjects (e.g. 150 years old). Truncating the simulated covariates to more realistic values could help mitigating this problem; however, this was not assessed in the present analysis primarily because the cut-off values should be selected on a case-by-case basis, whereas the present work aimed at maximizing the generalizability of the results. In terms of categorical covariates, no clear differences between the operating characteristics of the three methods stood out; poorly represented categories, like “Other” and “American Indian” for race, were characterized by large rBias and rRMSE (Figure 2C).

Because the relationship between some pairs of log-transformed covariates deviated significantly from linearity (see e.g. age∼WT correlation in Figure 1B), MVND provided more inaccurate and imprecise predictions of the covariance terms compared to BS and CD (Figure 3B-C). As expected, the BS could reproduce the original relationship between covariates regardless of its shape (Figure 3A), while CD appeared to be more robust than MVND to deviations from the linearity assumptions. It should be noted that the original data set was characterized by a substantial overlap in the covariate distributions between HS and patients (Figure 1); in case relevant differences between these distributions are observed, BS and MVND should account for that by using stratification and mixture modeling, respectively.

In Scenario B, the aim was to evaluate the extrapolation capabilities of MVND and CD by assessing their predictive performance on a test data set composed by subjects older than 55 years of age, when the simulations were informed by a training data set from younger subjects. Similar to Scenario A, the results suggested that in general CD achieved higher accuracy and precision levels than MVND for continuous covariates (Figure 4A-B). While comparable performance was observed for the prediction of the race categories, the predicted proportion of males and females was more biased and imprecise for CD compared to MVND. However, a previous study comparing MVND and CD approaches for multiple imputations found that CD is more accurate than MVND whenever the data include categorical variables [21]. Figure S2 indicates that - in the training data set - females have a generally lower WT than males, whereas in the test data set the WT distribution in male subjects is shifted towards that of females, and the two overlap into a unimodal distribution. It is reasonable to assume that, since WT is a relatively strong predictor for sex in the training data set, CD predicted a higher proportion of females (and lower proportion of males) because the mean WT in the test data set is lower than the one in the training data set. Instead, MVND led to better performance because the sex proportion is similar between the two data sets, though it should be noted that MVND can poorly perform when the marginal distributions are characterized by strong bimodalities [15].

When the HS data were removed from the training data set (Scenario C), the non-linearities in the correlation between age and other covariates were diminished (purple line in Figure 1B). As a result, the extrapolation performance of MVND improved compared to Scenario B and, although the CD operating characteristics also did, no large differences were observed between the two methods in terms of continuous covariates (Figure 4A-B). rBias and rRMSE in the sex covariate were still lower for MVND vs. CD, yet the aforementioned discussion on Scenario B concerning the different WT distribution between males and females applies to Scenario C as well (Figure S2).

In general, CD was more robust to departures of the data from normality assumptions. The reason seems to lie more on the PMM method used to predict the new covariate value rather than in the different statistical background between the two methods (in MVND the data are modeled as a sample from a joint multivariate normal distribution whereas in CD each variable is modeled conditionally on all the others). In fact, PMM slightly downweighs the model prediction by picking up a covariate value in the original data set that is very close to the predicted one; nevertheless, it should be noted that if strong non-linear relations exist in the original data set, the consequent misspecification in the underlying predictive model could result in poor CD performance [22]. Tannenbaum et al [15] suggest to carry out additional methods prior to creating the MVND in order to correct for non-linear relations between continuous covariates, but this was not performed in the present implementation (apart from log-transformation). Although this could have improved the MVND results, especially in terms of the variance-covariance matrix, it is likely that CD would have also benefitted from such pre-transformations given that it is still based on linear prediction models. Likewise, truncation of MVND simulations to plausible values might also have positively impacted the MVND operating characteristics. On the other hand, it can be noticed that while CD simulates ready-to-use covariates (Supplementary Information 2), MVND can require additional post-processing of its input as well as output data, which in turn requires extra efforts and assumptions to be made.

The different results of the MVND between Scenario B and C can be attributed primarily to the distribution of age in HS, which is approximately uniform between 18 and 45 years. As shown in Figure 1A, the HS data distort the overall distribution of age, which is instead well approximated by a normal distribution in patients alone. Presumably, this is also the reason for the MVND issues in reproducing dispersion metrics under Scenario A. This suggests that study-specific nuisance covariates, such as HS vs. patients, should be handled carefully before embarking in a CTS effort. For example, in order to mimic the data generation process, the MVND approach could have been applied separately for patients and HS; however, this stratification should be done carefully as it may decrease the precision in the means and covariances of the two MVNDs, thereby increasing the uncertainty in the corresponding predictions.

BS was excluded from Scenario B and C because, by definition, it does not allow to extrapolate outside of the empirical distribution. This limitation can be overcome if observational data on the target indication are available, for example from databases based on routine clinical care used to identify patients for enrollment in clinical trials. However, while for common indications like hypertension this might easily be achieved, large patient registries for less frequent diseases are often not available.

CD performed with the *mice* package requires the specification of some options such as the number of donors K or the type of prediction model to use. All the defaults options were used in this analysis, yet the sensitivity of the results to the options choice should be further investigated. An interesting feature of *mice* that was not explored in the present work is that it potentially enables simulation of time-varying covariates, which cannot be done with an MVND approach. Furthermore, extrapolation to other clinical contexts and generalizability of the results obtained in this work should be further explored.

The simulation of covariates, together with a disease and a drug model, represent the minimum working components that enable the generation of an in-silico response to treatment for a group of patients. Thus, besides providing a tool for prospective investigations of study design features, CTS represents an appealing methodology for the creation of synthetic controls in clinical trials. Synthetic controls are formally defined as statistical methods that can be used to evaluate the comparative effectiveness of an intervention using external control data [23]. Although synthetic controls are usually based on collected real evidence [24], the use of predictive statistical models to simulate virtual responses to standard-of-care treatments has recently been proposed as a way to generate synthetic controls [25]. In the case of rare diseases and small populations in general (e.g. pediatrics), where the use of control groups is often hampered by feasibility and ethical hurdles, simulated synthetic controls represent a promising alternative, as also indicated by their increased regulatory acceptance [26].

To conclude, the present analysis revealed that if CTS is used to simulate within the range of the observed distribution - for example when studies in the target population are already available from other sources - BS can be considered as the preferred method for covariates simulation, particularly because it is able to guarantee the physiological plausibility of the simulated covariates. On the other hand, if the CTS objectives involve extrapolation to new populations, a parametric method like CD or MVND is needed. In case the empirical multivariate distribution used to inform the extrapolation is characterized by linearly related covariates and unimodal marginal distributions, CD and MVND have comparable performance, and the use of MVND may be favored in light of its simpler statistical framework and well-established use. However, if uncertainty about the MVND hypotheses exists, CD allows to partly relax these hypotheses, thereby increasing the confidence in the simulation outcomes compared to MVND.

### Study Highlights

#### What is the current knowledge on the topic?

Common ways of generating virtual subjects to be used in clinical trial simulation (CTS) tasks are based upon bootstrap (BS) procedures or multivariate normal distributions (MVND).

#### What question did this study address?

What is the performance of an alternative method based on conditional distributions (CD) compared to BS and MVND?

#### What does this study add to our knowledge?

If CTS is used to simulate within the range of the observed distribution the BS is the preferred method for covariates simulation. When the CTS objectives involve extrapolations to new populations, a parametric method like CD or MVND is needed. As the MVND approach rests on relatively strong assumptions, i.e. linearly related covariates and unimodal distributions, CD is more robust when deviations from these assumptions occur.

#### How might this change drug discovery, development, and/or therapeutics?

Pharmacometricians will have at their disposal a new method for covariates simulation, which is particularly favorable when the empirical covariate distribution cannot be approximated by a MVND.

## Acknowledgments

We would like to thank all Pharmetheus’ colleagues for their valuable feedback on this work. Particular thanks go to Ola Caster, Sofia Friberg Hietala and Peter Milligan, whose critical input improved the quality of the manuscript.

## Author contributions

G.S. and E.N.J. wrote the manuscript; E.N.J. designed the research; G.S. and E.N.J. performed the research; G.S. analyzed the data.

## Supplementary information titles

Supplementary Information 1: Fully conditional specification (FCS) algorithm as implemented in the *mice* package.

Supplementary Information 2: R function developed for covariates simulation using CD with minimum working examples of use.

Figure S1: Illustrative example of the methodology used to simulate covariates with CD.

Figure S2: Histograms of the continuous covariates in the training and test data sets for Scenarios B and C, colored by sex.

