## Supplementary material for "Conditional distribution modeling as an alternative method for covariates simulation: comparison with joint multivariate normal and bootstrap techniques": Figure S1

**Figure S1**. Illustrative example depicting how FCS was used to simulate covariates with CD (only the first two iterations of FCS are shown), using a standard linear regression model instead of predictive mean matching as imputation model, under Scenario A. The starting point is represented by a dummy data set with the same rows as the original data set and all covariates assumed to be missing (green table), which is attached to the original data set identified by the orange color (0). At iteration 0 all the missing data are first imputed by drawing a random sample from the original data set on a covariate-by-covariate basis (1). During iteration 1, missing covariates are sequentially imputed: the covariate under question is modeled conditionally on all the others and the model prediction is used as imputation (2). When all the covariates have been imputed with the respective imputation model, a new full data set is obtained (3), which is then used as starting point for iteration 2. This iterative process goes on until convergence is reached, defined as the mean and standard deviation of each covariate being stable over the last iterations.

Legend: Original data set (orange rows); Simulated data set (green rows).

**Starting point**

| ID | AGE | WT | SCR | SEX | RACE |
|---|---|---|---|---|---|
| 1 | 24 | 84 | 122 | M | White |
| 2 | 42 | 64 | 53 | F | White |
| … | … | … | … | … | … |
| 939 | 44 | 83 | 106 | M | Asian |
| 1 | | | | | |
| 2 | | | | | |
| … | | | | | |
| 939 | | | | | |

Random draw of each covariate value from orginial data set

**Iteration 0, random draw**

| ID | AGE | WT | SCR | SEX | RACE |
|---|---|---|---|---|---|
| 1 | 24 | 84 | 122 | M | White |
| 2 | 42 | 64 | 53 | F | White |
| … | … | … | … | … | … |
| 939 | 44 | 83 | 106 | M | Asian |
| 1 | 42 | 84 | 122 | M | White |
| 2 | 20 | 64 | 53 | F | Black |
| … | … | … | … | … | … |
| 939 | 80 | 83 | 106 | M | White |

$$AGE = a_{AGE,0} + a_{AGE,1} \cdot WT + a_{AGE,2} \cdot SCR + a_{AGE,3} \cdot SEX + \sum_i a_{AGE,4i} \cdot RACE_i$$

**Iteration 1, Impute AGE**

| ID | AGE | WT | SCR | SEX | RACE |
|---|---|---|---|---|---|
| 1 | 24 | 84 | 122 | M | White |
| 2 | 42 | 64 | 53 | F | White |
| … | … | … | … | … | … |
| 939 | 44 | 83 | 106 | M | Asian |
| 1 | | 84 | 122 | M | White |
| 2 | | 64 | 53 | F | Black |
| … | | … | … | … | … |
| 939 | | 83 | 106 | M | White |

$$WT = a_{WT,0} + a_{WT,1} \cdot AGE + a_{WT,2} \cdot SCR + a_{WT,3} \cdot SEX + \sum_i a_{WT,4i} \cdot RACE_i$$

**Iteration 1, Impute WT**

| ID | AGE | WT | SCR | SEX | RACE |
|---|---|---|---|---|---|
| 1 | 24 | 84 | 122 | M | White |
| 2 | 42 | 64 | 53 | F | White |
| … | … | … | … | … | … |
| 939 | 44 | 83 | 106 | M | Asian |
| 1 | 44 | | 122 | M | White |
| 2 | 23 | | 53 | F | Black |
| … | … | | … | … | … |
| 939 | 70 | | 106 | M | White |

…

$$logit(RACE) = a_{RACE,0} + a_{RACE,1} \cdot AGE + a_{RACE,2} \cdot SCR + a_{RACE,3} \cdot SEX + a_{RACE,4} \cdot WT$$

**Iteration 1, Impute RACE**

| ID | AGE | WT | SCR | SEX | RACE |
|---|---|---|---|---|---|
| 1 | 24 | 84 | 122 | M | White |
| 2 | 42 | 64 | 53 | F | White |
| … | … | … | … | … | … |
| 939 | 44 | 83 | 106 | M | Asian |
| 1 | 44 | 80 | 115 | F | |
| 2 | 23 | 70 | 44 | M | |
| … | … | … | … | … | |
| 939 | 70 | 78 | 80 | M | |

**Iteration 2, full data set**

| ID | AGE | WT | SCR | SEX | RACE |
|---|---|---|---|---|---|
| 1 | 24 | 84 | 122 | M | White |
| 2 | 42 | 64 | 53 | F | White |
| … | … | … | … | … | … |
| 939 | 44 | 83 | 106 | M | Asian |
| 1 | 44 | 80 | 115 | F | Black |
| 2 | 23 | 70 | 44 | M | White |
| … | … | … | … | … | …. |
| 939 | 70 | 78 | 80 | M | White |

**Iteration 2, Impute AGE**

| ID | AGE | WT | SCR | SEX | RACE |
|---|---|---|---|---|---|
| 1 | 24 | 84 | 122 | M | White |
| 2 | 42 | 64 | 53 | F | White |
| … | … | … | … | … | … |
| 939 | 44 | 83 | 106 | M | Asian |
| 1 | | 80 | 115 | F | Black |
| 2 | | 70 | 44 | M | White |
| … | | … | … | … | …. |
| 939 | | 78 | 80 | M | White |

**Iteration 2, Impute WT**

| ID | AGE | WT | SCR | SEX | RACE |
|---|---|---|---|---|---|
| 1 | 24 | 84 | 122 | M | White |
| 2 | 42 | 64 | 53 | F | White |
| … | … | … | … | … | … |
| 939 | 44 | 83 | 106 | M | Asian |
| 1 | 43 | | 115 | F | Black |
| 2 | 22 | | 44 | M | White |
| … | … | | … | … | …. |
| 939 | 70 | | 80 | M | White |

…

**Iteration 2, Impute RACE**

| ID | AGE | WT | SCR | SEX | RACE |
|---|---|---|---|---|---|
| 1 | 24 | 84 | 122 | M | White |
| 2 | 42 | 64 | 53 | F | White |
| … | … | … | … | … | … |
| 939 | 44 | 83 | 106 | M | Asian |
| 1 | 44 | 81 | 110 | F | |
| 2 | 23 | 75 | 50 | M | |
| … | … | | … | … | |
| 939 | 70 | 69 | 88 | M | |
