## Supplementary information 1 for "Conditional distribution modeling as an alternative method for covariates simulation: comparison with joint multivariate normal and bootstrap techniques"

**FCS algorithm**

1. Specify an imputation model $P(Y_j^{mis}|Y_j^{obs}, Y_{-j}, \phi)$ for variable $Y_j$ with j=1,…,p, where p is the number of variables
2. For each j, fill in starting imputations $\dot{Y}_j^0$ by random draws from $Y_j^{obs}$
3. Repeat for t=1, …, M, where M is the number of iterations
4. Repeat for j=1, …, p
5. Define $\dot{Y}_{-j}^t = (\dot{Y}_1^t, \dots, \dot{Y}_{j-1}^t, \dot{Y}_{j+1}^{t-1}, \dots, \dot{Y}_p^{t-1})$ as the currently complete data except $Y_j$
6. Draw $\dot{\varphi}_j^t \sim P(\dot{\varphi}_j^t|Y_j^{obs}, \dot{Y}_{-j}^t)$
7. Draw imputations $\dot{Y}_j^t \sim P(Y_j^{mis}|Y_j^{obs}, \dot{Y}_{-j}^t, \dot{\varphi}_j^t)$
8. End repeat j
9. End repeat t

**Definitions**

- Y: matrix with p columns representing the data set with missingness
- $Y_j^{mis}$: set of missing values for the j-th variable
- $Y_j^{obs}$ : set of observed values for the j-th variable
- $Y_{-j}$: values of all variables in Y that are not the j-th one
- $\phi$: imputation model with parameters φ
