## Supplementary information 2 for "Conditional distribution modeling as an alternative method for covariates simulation: comparison with joint multivariate normal and bootstrap techniques"

##------------------------------------------------------------------------------------
##          File name: simCovMICE.R
##          Created: 2021-02-15
##          Use: function creating covariates distribution using mice
##          Required packages: 
##               - 'dplyr'  version 0.8.3  
##               - 'mice'   version 3.6.0  
##
##          By: Giovanni Smania
##------------------------------------------------------------------------------------

simCovMICE <- function(m = 5, 
                       orgCovs, 
                       catCovs = c("SEX","RACE"), 
                       seedCovs = NULL,
                       targetRangeSeedCovs = NULL,
                       seedCovsValues = NULL,
                       nsubj = nrow(orgCovs),
                       contMeth = "pmm",
                       sampleFromReal = T,
                       ...)
  #####
## INPUTS:
##    m        --> how many replicates of the original data to generate
##    orgCovs  --> n x p dataframe containing the original, observed, time-invariant covariates (ID should not be included) that will be used to inform the imputation
##    catCovs  --> character vector containing the name of the categorical covariates in orgCovs
##    seedCovs --> character vector containing the name of the covariates that should be used to seed the imputation, they will be sampled from the original data set
##    targetRangeSeedCovs --> if the seeding is based on a desired range, then define it here
##    seedCovsValues --> vector of length n containing the seeding cov values 
##    nsubj   --> number of simulated subjects, default is subjects orgCovs
##    contMeth   --> method used to predict continuous covariates within mice, default is 'pmm'
##    sampleFromReal --> should seedCovs be sampled from orgCovs?
##    ... --> additional input to mice call
##
## OUPUT: a data frame with the simulated covariates, with nsubj*m rows and (p+1) columns
## 
## NOTES: missing values in orgCovs must be coded as NA
#####
{
  
  # names of continuous covariates
  contCovs <- setdiff(names(orgCovs),catCovs)
  
  # create copy of the original data set with factor version of categorical covariates
  orgCovsF <- orgCovs %>% dplyr::mutate_at(catCovs,function(x) as.factor(x))
  
  ## find covariates with missing values
  missVars <- names(orgCovs)[colSums(is.na(orgCovs)) > 0]
  
  # impute missing data once with mice  
  if(length(missVars)>0)
  {
    imp1 <- mice::mice(orgCovsF, m=1, printFlag=FALSE, maxit = 15)
    orgCovs <- mice::complete(imp1)
  }
  
  miCovs <- orgCovs[1:nsubj,] %>% mutate_all(function(x) NA)
  
  
  if(!is.null(seedCovs)) 
  {
    if(sampleFromReal)
    {
      ## create vector of seedCov values contained in original data set
      poolSeed <-  orgCovs[orgCovs[,seedCovs]>=min(targetRangeSeedCovs) & orgCovs[,seedCovs]<=max(targetRangeSeedCovs),seedCovs]
      ## do the actual sampling
      miCovs[seedCovs] <- sample(poolSeed,nsubj,replace = T)
    }
    else
      miCovs[seedCovs] <-  seedCovsValues 
  }
  
  combCovs <- orgCovs %>% mutate(Type="Original") %>% 
    bind_rows(miCovs %>% mutate(Type="Simulated")) %>%
    mutate_at(catCovs,function(x) as.factor(x))
  
  myPredMat <- mice::make.predictorMatrix(combCovs)
  myPredMat[,c("Type")] <- 0
  
  myMethods <- mice::make.method(combCovs)
  myMethods[contCovs] <- contMeth
  
  imp2 <-mice::mice(combCovs, m=m,printFlag=FALSE,predictorMatrix = myPredMat, method = myMethods, ...)
  
  impCovs <- mice::complete(imp2,action="long") %>% 
    filter(Type=="Simulated") %>% 
    mutate(NSIM=.imp) %>% 
    select(everything(),-.id,-Type,-.imp)
  
  return(impCovs)
  
}


##------------------------------------------------------------------------------------
##
##      MINIMUM WORKING EXAMPLE ON THE USE OF simCovMICE()
##
##------------------------------------------------------------------------------------

library(dplyr)

# A) Simulations without seeding (all covariates are simulated)

orgCovsEx <- data.frame(AGE = c(50,42,39,70),
                        WT  = c(84,64,88,55),
                        SMOKE = c(1,2,2,1))

myCovSimMICE_A <- simCovMICE(m = 2,orgCovs = orgCovsEx,catCovs = c("SMOKE"))
myCovSimMICE_A

# B) Simulations using AGE as a seeding covariate: AGE is assumed to be 
#    known, and sampled from the original data set, using only values within 40 and 50 years.

myCovSimMICE_B <- simCovMICE(m = 2,orgCovs = orgCovsEx,catCovs = c("SMOKE"),seedCovs = "AGE",
                             sampleFromReal = T,targetRangeSeedCovs = c(40,50))

myCovSimMICE_B

# C) Simulations using AGE as a seeding covariate: AGE is assumed to be 
#    known, and equal to the age vector obtained in the first simulated data set of example A

myCovSimMICE_C <- simCovMICE(m = 2,orgCovs = orgCovsEx,catCovs = c("SMOKE"),seedCovs = "AGE",
                             sampleFromReal = F,seedCovsValues = myCovSimMICE_A$AGE[myCovSimMICE_A$NSIM==1])

myCovSimMICE_C
