## Supplementary figures and images for "Conditional distribution modeling as an alternative method for covariates simulation: comparison with joint multivariate normal and bootstrap techniques"

### Figure S2

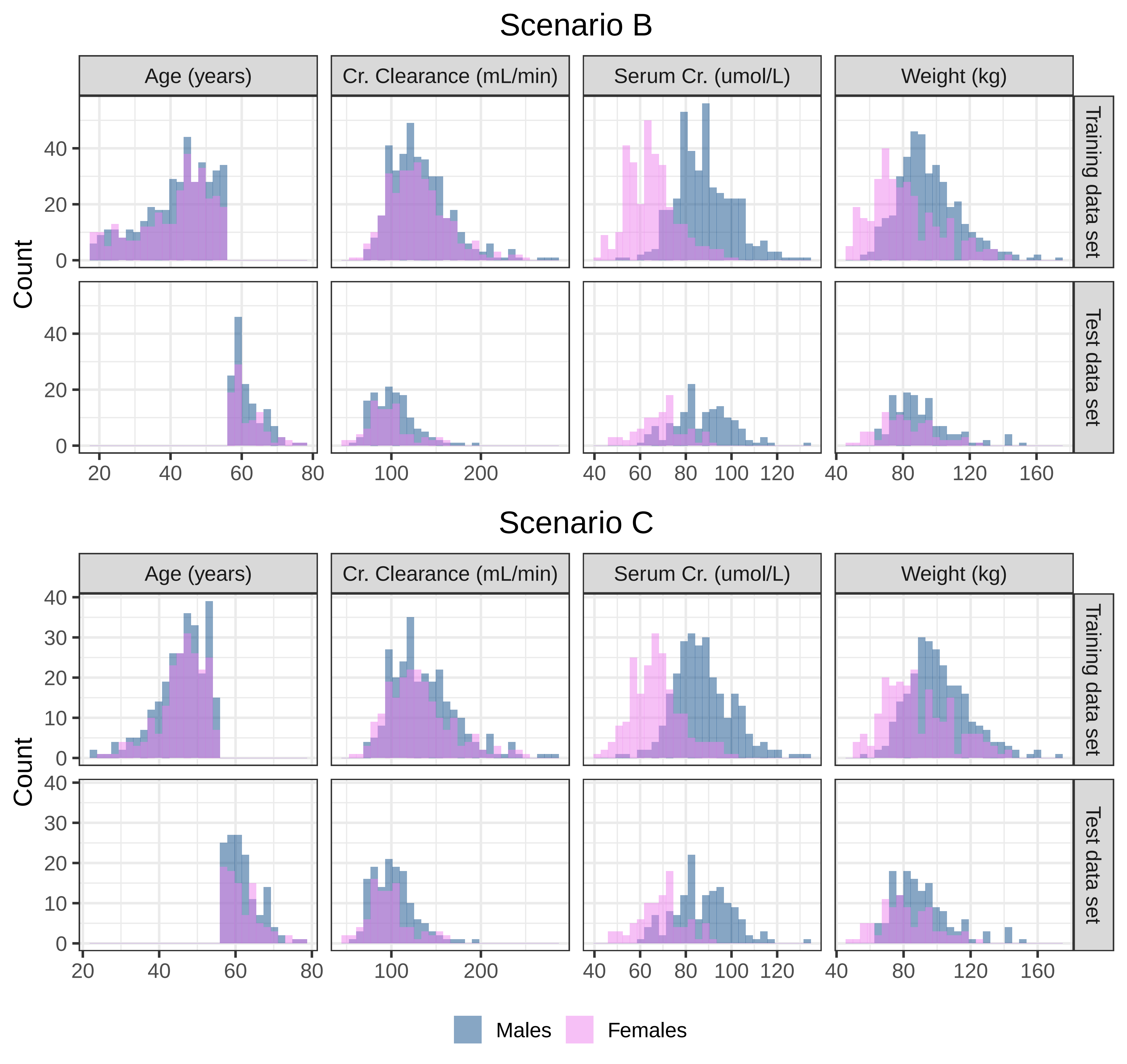
